# Adaptor linked K63 di-Ubiquitin activates Nedd4/Rsp5 E3 ligase

**DOI:** 10.1101/2021.11.25.470069

**Authors:** Lu Zhu, Qing Zhang, Ciro Cordeiro, Sudeep Banjade, Richa Sardana, Yuxin Mao, Scott D. Emr

## Abstract

Nedd4/Rsp5 family E3 ligases mediate numerous cellular processes, many of which require the E3 ligase to interact with PY-motif containing adaptor proteins. Several Arrestin-Related Trafficking adaptors(ARTs) of Rsp5 were self-ubiquitinated for activation, but the regulation mechanism remains elusive. Remarkably, we demonstrate that Art1, Art4, and Art5 undergo K63-linked di-ubiquitination by Rsp5. This modification enhances the PM recruitment of Rsp5 by Art1 or Art5 upon substrate induction, required for cargo protein ubiquitination. In agreement with these observations, we find that di-ubiquitin strengthens the interaction between the Pombe orthologs of Rsp5 and Art1, Pub1 and Any1. Further, we discover that the HECT domain exosite protects the K63-linked di-ubiquitin on the adaptors from cleavage by the deubiquitination enzyme Ubp2. Strikingly, loss of this protection results in the loss of K63-linked di-ubiquitin from the adaptors and diverts the adaptors for K48-linked poly-ubiquitination and proteasome-mediated degradation. Together, our study uncovers a novel ubiquitination modification implemented by Rsp5 adaptor proteins, underscoring the regulatory mechanism of how adaptor proteins control the recruitment and activity of Rsp5 for the turnover of membrane proteins.

## Introduction

The Nedd4/Rsp5 family E3 ligases are responsible for membrane protein ubiquitination, required for endocytosis and lysosome-dependent protein degradation. Tryptophan-tryptophan (WW) domains of Nedd4 family E3 ligases bind to substrate proteins via interaction with PY motifs containing a consensus sequence P/L-P-x-Y (Rotin & Kumar, 2009; Schild *et al*, 1996). Other substrates lack PY motifs and instead rely on interactions with adaptor proteins that recruit the Nedd4 E3 ligase to them, exemplified by a family of arrestin-related trafficking adaptors (ARTs) that bridge the association between substrates and Rsp5 for ubiquitination(Lin *et al*, 2008). Additionally, Rsp5 adaptors include a diverse group of transmembrane (TM) proteins to mediate degradation of membrane proteins localized at the PM, Golgi, endosome and vacuole membrane (Alvaro *et al*, 2014; Becuwe *et al*, 2012; Hatakeyama *et al*, 2010; Hettema *et al*, 2004; Hovsepian *et al*, 2018; Leon *et al*, 2008; Li *et al*, 2015; MacDonald *et al*, 2012; Nikko & Pelham, 2009; O’Donnell *et al*, 2013; Sardana *et al*, 2018; Zhu *et al*, 2020)

Many of the Nedd4/Rsp5 adaptor proteins undergo self-ubiquitination. The ART proteins Art1, Art4 and Art8 require specific ubiquitination by Rsp5 to reach full activity (Becuwe *et al*., 2012; Hovsepian *et al*, 2017; Lin *et al*., 2008). Ubiquitination of Nedd4 adaptor protein *Commissureless* is required to downregulate the Robo receptor at the cell surface of axons, essential for midline crossing (Ing *et al*, 2007; Myat *et al*, 2002). The N-lobe region of the Nedd4/Rsp5 family E3 ligase HECT domain contains an exosite which binds ubiquitin and has been shown to orient the ubiquitin chain to promote conjugation of the next ubiquitin molecule of the growing polyubiquitin chain(Kim *et al*, 2011; Maspero *et al*, 2011). It was proposed that ubiquitinated Rsp5 adaptors are more active when locked onto Rsp5 but less active when unlocked by Ubp2 (MacDonald *et al*, 2020). However, the mechanism of how Nedd4/Rsp5 adaptor ubiquitination helps enhance E3 ligase function remains unclear.

In this study, we decoded the activation mechanism of how adaptor protein ubiquitination enhances E3 ligase function and how this ubiquitination itself is regulated by the deubiquitination (DUB) enzyme Ubp2. Remarkably, we discovered that the Rsp5 adaptors Art1, Art4, and Art5 are conjugated with K63-linked di-Ub at specific ubiquitination sites. Ubiquitination of Art5 and Art1 enhances Rsp5 recruitment to the plasma membrane thereby promoting substrate ubiquitination. Our analysis of the binding affinity of di-Ub or isolated PY motifs to Rsp5 targeted domains uncovered that K63-linked di-Ub conjugation to the adaptor protein Any1 sharply enhances its binding to E3 ligase Pub1. Strikingly, we found that deletion of *UBP2* rescues the deubiquitination of adaptor proteins Art5 and Art1 in the *rsp5*-exosite mutant. Our data reveals the interplay between Ubp2 and “Rsp5 exosite engagement” to modulate adaptor protein ubiquitination and catalyze the switch from K63-linked di-Ub to K48-linked ubiquitin chains. Taken together, these results serve as a portal for future studies of Nedd4/Rsp5 adaptor proteins in general.

## Results

### Rsp5 adaptor protein Art5 undergoes K63-linked di-ubiquitination

In yeast, 14 α-arrestin domain containing proteins have been identified: Art1-Art10(Lin *et al*., 2008; Nikko & Pelham, 2009), Bul1-Bul3 (Yashiroda *et al*, 1996) and Spo23 (Aubry & Klein, 2013). These proteins have clear arrestin sequence signatures and contain multiple PY motifs that specifically interact with the WW domains in Rsp5 (Baile *et al*, 2019), and can recruit Rsp5 to specific intracellular locations. This interaction not only results in ubiquitination of cargo proteins, but also ubiquitination of ARTs themselves. In fact, several α-arrestin domain containing proteins have been shown to be ubiquitinated by Rsp5, including Bul1, Bul2, Art1, Art4, Art5, Art6 and Art8. Among these, Art5 contains an α-arrestin domain and three C-terminal PY motifs (figure 1A). It has been shown that Art5 is the only ART protein required for the inositol-induced endocytosis and degradation of the plasma membrane (PM) inositol transporter Itr1 (Nikko & Pelham, 2009).

**Figure 1.**
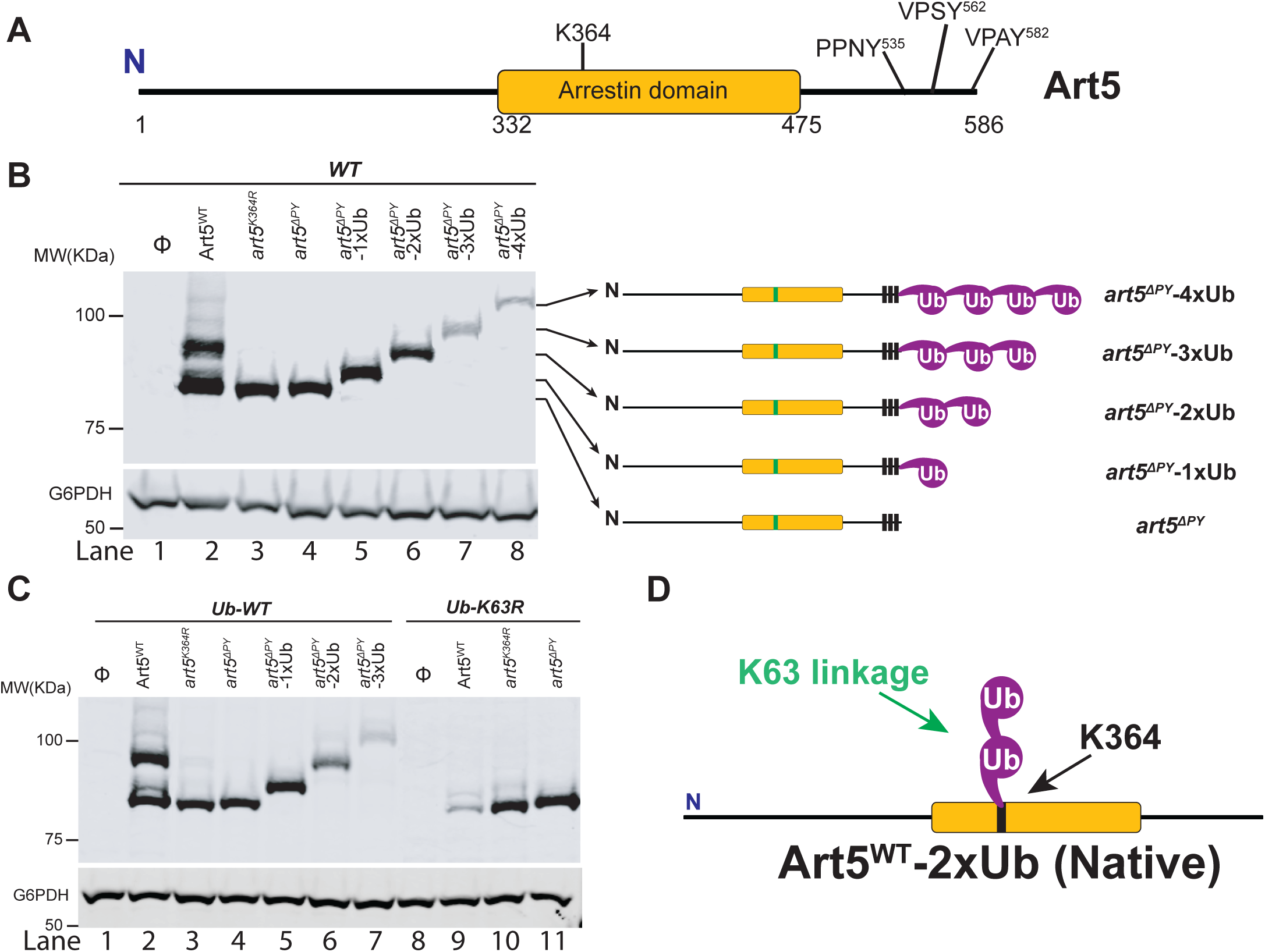
Art5 undergoes K63-linked di-ubiquitination. (A) Schematic representation of the domain architecture of Art5. (B) A di-ubiquitin is conjugated at K364 residue of Art5. Western blot analysis of Art5, *art5^K364R^*, *art5^ΔPY^*, *art5^ΔPY^*-1xUb, *art5^ΔPY^*-2xUb, *art5^ΔPY^*-3xUb and *art5^ΔPY^*-4xUb in the wild-type strain. (C) Art5 is di-ubiquitinated in a K63 linkage at the residue K364. Western blot analysis of Art5, *art5^K364R^*, art*5^ΔPY^* in both the *Ub-WT* and *Ub-K63R* mutant strains. (D) Model depicting the K63-linked di-ubiquitination of Art5 at the K364. The whole cell lysate protein samples were resolved on 7% SDS-PAGE gels and the blot was probed with FLAG and GAPDH antibodies.

We found that at steady state, endogenous Art5 migrates in two major bands by SDS-PAGE, corresponding to the ubiquitinated and non-ubiquitinated species (Lane 2 in the Fig. 1B). Mass spectrometry has previously indicated that ubiquitin is mainly conjugated on the K-364 residue of the Art5 α-arrestin domain (Swaney *et al*, 2013). We confirmed that Art5 ubiquitination was nearly completely ablated by mutating K364(Fig. 1B, lane 3), and is completely abolished in the *art5*^ΔPY^ mutant in which all three PY motifs were mutated (lane 4), demonstrating that Art5 ubiquitination depends on its interaction with Rsp5 via PY motifs. There is a minor portion (albeit weak) of PY motifs dependent Art5 higher molecular weight species (lane 2), probably due to other lysines. Strikingly, the molecular weight difference (∼20KDa) between the non-ubiquitinated and ubiquitinated forms of Art5 appears to be more than one single ubiquitin (∼9KDa), suggesting more than one ubiquitin molecule is conjugated to the Art5 protein. To test this hypothesis, we fused the C-terminus of *art5*^ΔPY^ with 1, 2, 3 or 4 ubiquitin molecules to create *art5*^ΔPY^-1xUb, *art5*^ΔPY^-2xUb, *art5*^ΔPY^-3xUb and *art5*^ΔPY^-4xUb, respectively. Remarkably, the ubiquitinated Art5^WT^ runs in line with *art5*^ΔPY^-2xUb, indicating that Art5 is di-ubiquitinated mainly at the K-364 residue (Lane 4, 5 and 6 of Fig. 1B).

We next asked what is the linkage in the di-ubiquitin that is conjugated to Art5. Rsp5 mainly catalyzed K63-linked ubiquitin chain synthesis *in vivo* and *in vitro* (Lauwers *et al*, 2009; Saeki *et al*, 2009). We therefore decided to examine whether the di-ubiquitin moiety on Art5 is K63-linked. We analyzed the migration of Art5^WT^*, art5*^K364R^, and *art5*^ΔPY^ proteins in yeast strains expressing Ub-WT and Ub-K63R. Notably, as seen in Figure 1C, we found that the size of the di-ubiquitinated Art5 band (lane 2 and 3) is reduced to the mono-ubiquitinated band (lane 9 and 10), in line with *art5*^ΔPY^-1xUb. As expected, this mono-Ub was conjugated to K364 residue. We noticed the loss of Art5 protein in the Ub-K63R mutant in a K364 and PY motif dependent manner (lane 9, figure 1c), which will be discussed later. In addition, the mono-ubiquitinated band of Art5 in the yeast Ub-K63R mutant is K364 residue dependent, confirming that the mono-ubiquitin is conjugated mainly at the K364 residue. These data are consistent with the alignment with *art5*^ΔPY^ -2xUb? (Fig. 1C), indicating that endogenous Art5 is di-ubiquitinated. Together, our result demonstrate that Art5 protein is di-Ubiquitinated at residue K364 in a K63 linkage by Rsp5.

Besides Art5, we next addressed if other ART proteins also undergo K63-linked di-ubiquitination. To test this idea, we employed the same approach to analyze another ART family member, Art1. Art1 was found to mediate downregulation of plasma membrane nutrient transporters such as Can1, Mup1, Fur4, and Lyp1. Art1 contains an N-terminal arrestin fold with PY motifs near its C-terminus (Fig. S1A), which bind to Rsp5’s WW domains. The K486 residue is required for Art1 ubiquitination (Lin et al., 2008). As anticipated, the ubiquitinated form of Art1 shows the same mobility shift in comparison with *art1*^ΔPY^-2xUb (Fig. S1B). To test if Art1 is ubiquitinated in a K63 linkage, we expressed the Art1^WT^, *art1*^K486R^ and *art1*^ΔPY^ in a yeast strain expressing only Ub-K63R. The ubiquitinated band of Art1 migrates with *art1*^ΔPY^-2xUb in the Ub-WT strain, while Art1 is mono-ubiquitinated at K486 in the yeast strain bearing Ub-K63R (Fig. S1C).

In addition to Art5 and Art1, another α-arrestin domain containing protein, Art4, also interacts with Rsp5 via PY motifs and can be ubiquitinated at a cluster of lysines (K235, K245, K264 and K267) in the N-terminal arrestin domain (Becuwe et al., 2012), as shown in Figure S1E. To examine the Art4 ubiquitination status, we expressed the Art4^WT^, *art4*^4KR^ and *art4*^ΔPY^ proteins in the yeast strains expressing Ub-WT and Ub-K63R. Due to Art4 phosphorylation when cells were grown in lactate medium, Art4 protein lysates were treated with phosphatase after being shifted to glucose containing culture medium. The ubiquitinated form of Art4^WT^ migrates with the *art4*^ΔPY^-2xUb (Fig. S1F). In contrast, Art4^WT^ was only mono-ubiquitinated in Ub-K63R condition. Taken together, our results demonstrated that α-arrestin domain containing adaptor proteins Art1, Art4 and Art5 are di-ubiquitinated and the di-ubiquitin is K63 linked (Fig. 1D, S1D and S1G).

### Ubiquitination of Art5 is required for cargo protein Itr1 ubiquitination

As shown in the figure 1B, Art5 ubiquitination depends on the interaction with Rsp5 and is abrogated in the Art5 K364R mutant. We therefore sought to investigate how Art5 ubiquitination affects efficient inositol-dependent endocytosis and protein degradation of Itr1. To determine whether Art5 ubiquitination is important for Art5 function, we expressed Art5^WT^ and *art5*^K364R^ in an *art5*Δ mutant bearing a chromosomal Itr1-GFP. Itr1-GFP degradation occurs after treatment with inositol in a dose-dependent manner. Thus, higher inositol concentrations applied for the same amount of time results in more Itr1-GFP degradation in the WT cells (Fig. 2A, lane 5-8; Fig. 2C) and protein sorting into the vacuole lumen (Fig. 2B, middle panels), and this endocytosis and degradation is Art5 dependent(Fig.2A, lane 1-4; Fig. 2C) (Nikko and Pelham, 2009). Cells expressing *art5*^K364R^ caused a severe decrease in the rate of Itr1-GFP degradation (Fig. 2A, lane 9-12; Fig. 2C) and protein endocytosis (Fig. 2B, right) at higher inositol concentrations compared with Art5-WT. Thus, Art5 ubiquitination is essential to promote efficient Itr1 endocytosis and protein degradation upon inositol-treatment.

**Figure 2.**
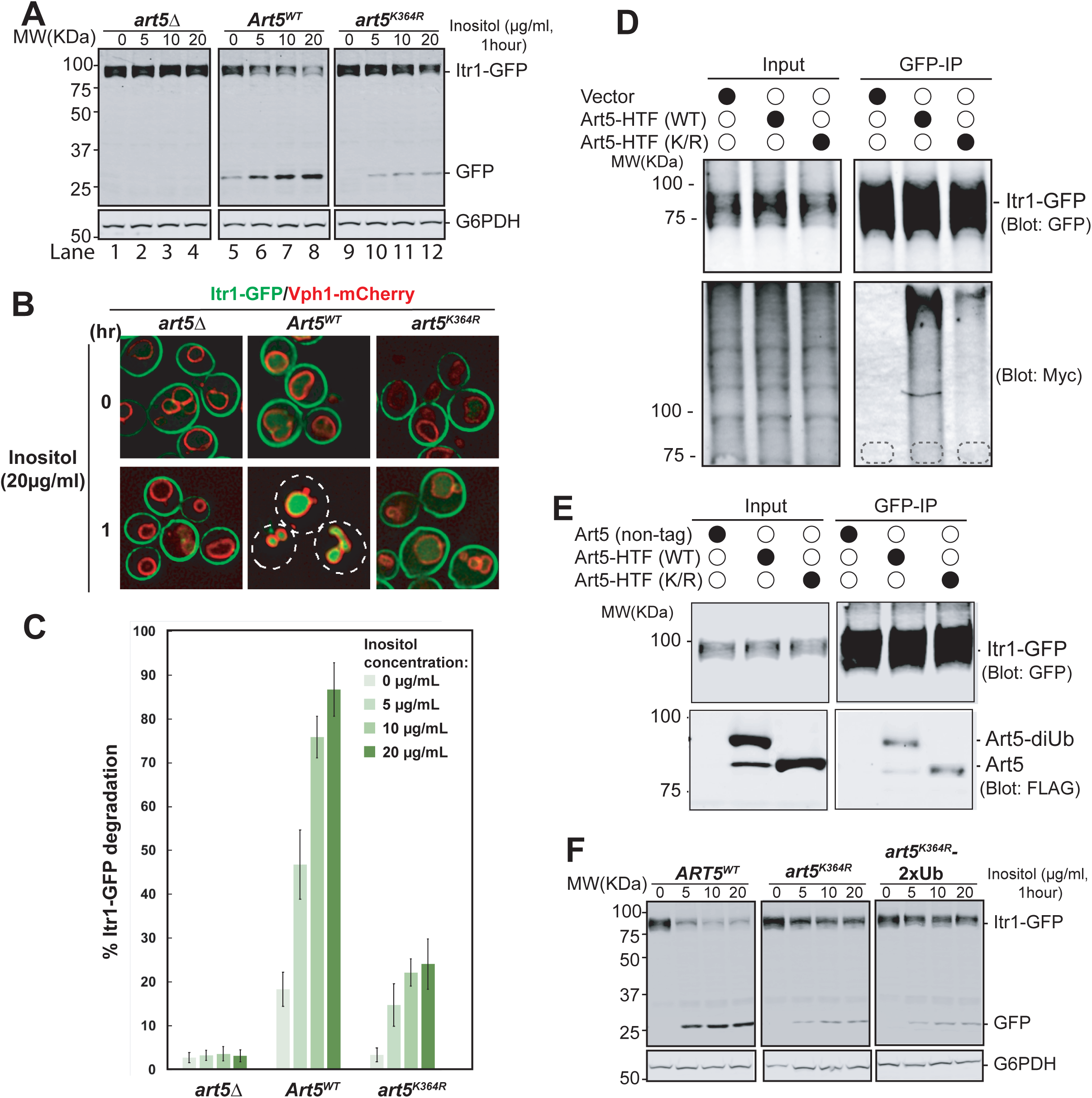
Ubiquitinated Art5 promotes cargo protein Itr1 ubiquitination. (A) Immunoblot analysis of Itr1-GFP endocytosis induced with indicated concentration of inositol for 60 minutes. (B) Fluorescence microscopy of *art5Δ*, Art5^WT^ or *art5^K364R^* cells expressing Itr1-GFP and vacuole membrane marker Vph1-mCherry with or without inducing endocytosis by treating with serial dilution of inositol. (C) Band densities of blots in (A) were quantified and expressed as the mean% Itr1-GFP degradation. *p*<0.001, n=3. (D) *doa4Δpep4Δart5Δ* cells expressing Itr1-GFP and Art5^WT^ or *art5^K364R^* were grown to mid-log phase in synthetic medium at 30°C. Cells were pretreated with 0.1µM CuCl2 for 4 hours to induce the Myc-Ub expression before treated with 20µg/ml of inositol. Cells were collected before and after 15 minutes of inositol treatment. Itr1-GFP was immunoprecipitated by GFP-Trap nanobody resin. Whole cell lysate and the IP reaction was resolved on 10% SDS-PAGE gels and the blots were probed with both GFP and Myc antibodies. The empty strain (*doa4Δpep4Δart5Δ*) is used as a negative control here. The whole cell lysate proteins in the left gels represent the loading control and the co-immunoprecipitated protein samples were resolved in right gels. (E) IP of Itr1-GFP and blotting for Art5^WT^ or *art5^K364R^*.

We hypothesize that the Itr1 sorting defect in *art5*^K364R^ is due to defective Itr1 ubiquitination. To test it, we expressed Itr1-GFP in a *doa4*Δ mutant bearing a Myc-Ub expression vector to stabilize ubiquitinated membrane proteins after multivesicular body sorting into the vacuole. After inositol treatment, Itr1-GFP was immunoprecipitated from cell lysates prepared from yeast expressing Art5^WT^ and *art5*^K364R^. The ubiquitinated pool of Itr1-GFP can be detected in the Art5^WT^ condition, whereas this ubiquitination was attenuated in *art5*^K364R^ condition (Fig. 2D). We next asked if the ubiquitination defect of Itr1 is due to the loss of protein-protein interaction between *art5*^K364R^ and Itr1. To test this, Itr1-GFP was immunoprecipitated from yeast strains expressing Art5^WT^ or *art5*^K364R^ (Fig. 2E). *art5*^K364R^ can be co-immunoprecipitated by Itr1-GFP comparable to Art5^WT^, indicating that the decrease of Itrt1 ubiquitination upon inositol stimulation is not due to the loss of interaction between adaptor protein Art5 and cargo protein Itr1.

The importance of ubiquitination of the ART proteins in cargo protein sorting is further supported by our previous findings that the *art1*^K486R^ allele results in a canavanine hypersensitivity phenotype (Lin et al., 2008). Here, we set out to test the endocytosis and protein degradation of Mup1-GFP in Art1^WT^ and *art1*^K486R^ after treatment with increased methionine concentrations. Consistent with previous results, the *art1*^K486R^ allele leads to a sorting defect of Mup1-GFP (Fig. S2A, S2B and S2C). Similarly, we sought to test if the Mup1-GFP can bind to both Art1^WT^ and *art1*^K486R^. To do so, we examined the protein interaction between Mup1 and Art1 using co-IP analysis. Indeed, we can observe the interaction between Mup1 and overexpressed Art1 (Fig. S2D). In agreement with previous finding that the acidic patch in the Mup1 N-terminal tail is required for binding with Art1 (Guiney *et al*, 2016), we showed that the Q49R Mup1 mutant did not interact with Art1 (Fig. S2E). Further, both Art1^WT^ and *art1*^K486R^ can bind to Mup1, as evidenced by the Co-IP of *art1*^K486R^ with Mup1 when Art1 ubiquitination is impaired (Fig. S2F). Thus, our results demonstrate that the sorting defect of Mup1-GFP in the presence of *art1*^K486R^ is not due to the loss of protein interaction between the adaptor protein Art1 and cargo protein Mup1.

Since TORC1 kinase regulates the Art1-dependent ubiquitin-mediated cargo protein endocytosis by modulating Art1 phosphorylation via Npr1 kinase (MacGurn *et al*, 2011), we decided to test if the non-ubiquitinated pool of Art1 loses the Npr1 dependence for phosphorylation thereby affecting cargo protein sorting. First, we expressed Art1^WT^ or *art1*^K486R^ in WT and *npr1*Δ mutant strains. We observed that both the di-ubiquitinated Art1 or the non-ubiquitinated Art1 pools migrated slightly faster in the *npr1*Δ mutant, consistent with dephosphorylation (Fig. S2G). Next, we treated the cells with either rapamycin or cycloheximide to monitor the change in phosphorylation status for ubiquitinated or non-ubiquitinated Art1. As shown in the figure S2H, the activated Npr1 kinase triggered by rapamycin treatment leads to phosphorylation of both Art1^WT^ and *art1*^K486R^; whereas the dephosphorylation of these two proteins is observed following cycloheximide treatment. The Npr1 kinase-dependent phosphorylation is therefore the intrinsic feature of Art1, regardless of the ubiquitination status of Art1.

Since ARTs ubiquitination enhances function, we next sought to test if C-terminal fusion with ubiquitin molecules could rescue the cargo sorting defect of *art1*^K486R^ or *art1*^ΔPY^. Since toxic arginine analog canavanine is transported by PM transporter Can1 in yeast and Can1 endocytosis prevents subsequent cell death (Grenson *et al*, 1966), canavanine hypersensitivity occurs when Can1 cannot be endocytosed (such as in an *art1Δ* mutant), which provides a readout of Art1 function. Thus, we examined the canavanine sensitivity of the *art1Δ* mutant expressing Art1^WT^, *art1*^K486R^ or *art1*^ΔPY^ fused with C-terminal 1x, 2x or 3xUb. The C-terminal fusions with ubiquitin molecules did not enhance the functionality of Art1^WT^, *art1*^K486R^ or *art1*^ΔPY^ (Fig. S2I, S2J). Besides Art1, we tested if Itr1-GFP sorting can be restored by *art5*^K364R^ with C-terminal 1xUb or 2xUb and found the 1x or 2xUb fusions do not enhance the Itr1 sorting (Fig. 2F). Together, our data indicate that di-ubiquitin needs to be conjugated at specific residues for proper functionality.

### PM recruitment of Rsp5 is enhanced by Art5 and Art1 protein ubiquitination

The *art5*^K364R^ mutant partially blocks the ubiquitination and cargo sorting of Itr1 after inositol treatment (Figure 2A and 2D), but still interacts with Itr1. We therefore hypothesize that the defective ubiquitination of *art5*^K364R^ may impair Rsp5 recruitment to the PM. To test this idea, we examined the localization of Art5-GFP in yeast cells before and after inositol treatment. As seen in figure 3A, the Art5^WT^-GFP localized at cytosol, nucleus and occasional cytosolic puncta (Sec7-negative, Figure 3A). Strikingly, the Art5^WT^-GFP is re-localized to PM puncta and patch structures after 1 hour of inositol (20µg/ml) treatment. This result is in line with our previous finding that Art5 is partially translocated to peripheral puncta after shift from minimal media to YPD (Lin *et al*., 2008), probably because the inositol concentration in YPD is not high enough to drive significant Art5 re-localization and Itr1 endocytosis. In comparison to Art5^WT^, the *art5*^K364R^-GFP and *art5*^ΔPY^ mainly remain in the cytosol even after inositol treatment (Figure 3B-D). We conclude that ubiquitination of Art5 is important for protein re-localization to the PM upon inositol treatment. We next asked if Rsp5 can be re-localized to the PM in an Art5-dependent manner after adding inositol to the growth media. As expected, Rsp5 was observed to be recruited to PM patches after inositol treatment in WT cells. However, Rsp5 PM recruitment after inositol treatment was substantially reduced in cells expressing either *art5*^K364R^ or *art5*^ΔPY^ (Figure 3E and 3F).

**Figure 3.**
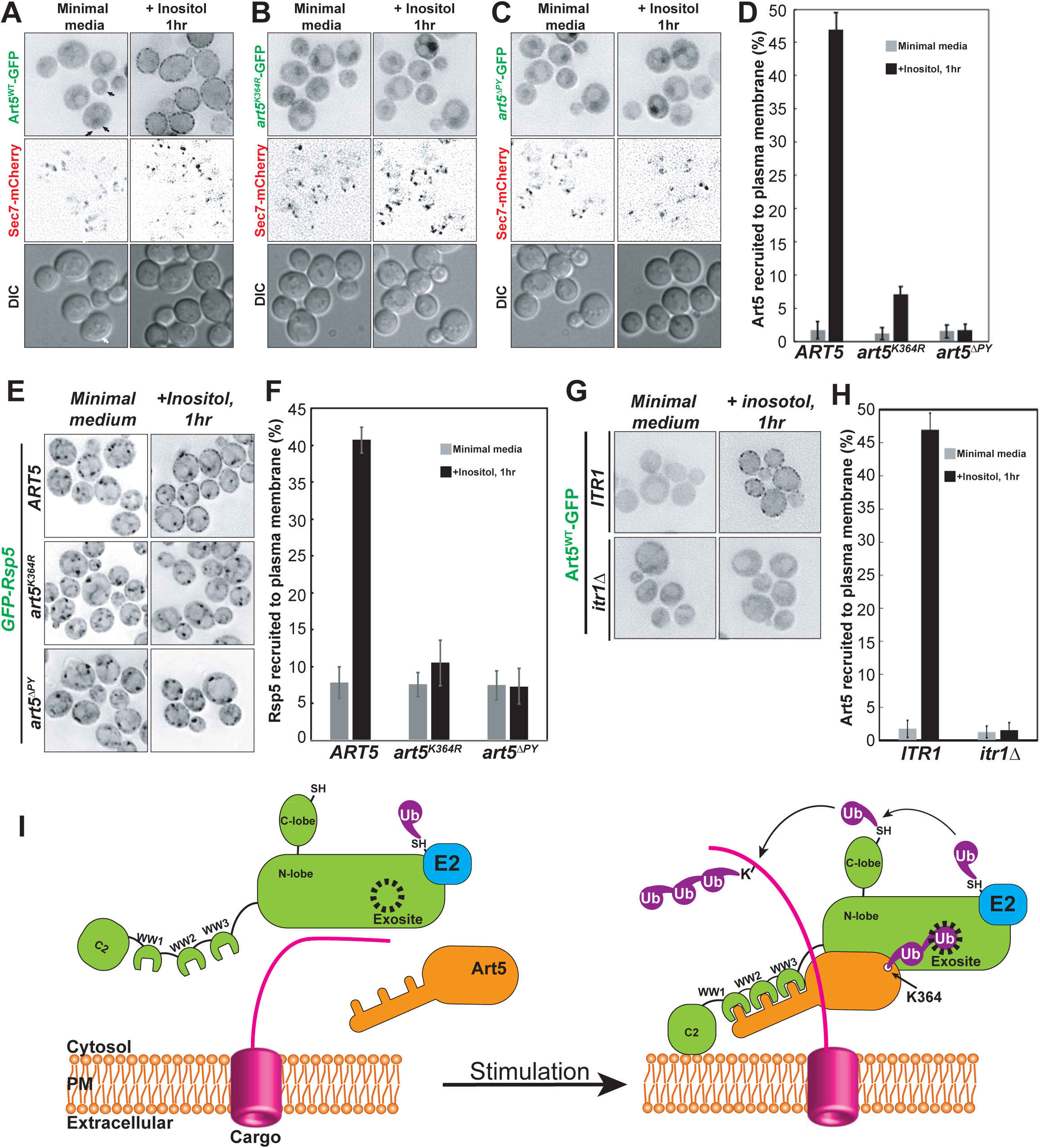
Rsp5 PM recruitment is enhanced by Art5 ubiquitination. (A-C) Fluorescence microscopy of Art5-GFP PY motif and K364R mutants in minimal media and after treating with inositol for 60min. (D) Quantification of PM localization of the indicated Art5 PY motif and K364R mutants. (E) Localization of GFP-Rsp5 in the presence of *ART5-WT*, PY motif or K364R mutants before and after inositol treatment for 60min. (F) Quantification of PM localization of Rsp5 in the experiment (E). (G-H) Fluorescence microscopy and quantification analysis of Art5-GFP in the WT and *itr1*Δ mutant condition, before and after inositol treatment. Scale bar = 2µm.

In addition to Art5, we also examined the PM localization of Art1 upon methionine treatment. Our previous results showed that Art1 is localized to Golgi, PM and cytosol, whereas the *art1*^K486R^ mutant is mainly localized to the cytosol(Baile *et al*., 2019; Lin *et al*., 2008). Art1 is recruited to the PM during cargo downregulation upon cycloheximide treatment or shift from synthetic medium to rich medium (Lin *et al*., 2008). As seen in the figure S3A, Art1 is efficiently recruited to the PM in YPD or in methionine media. In contrast to Art1^WT^, the recruitment of *art1*^K486R^ to the PM is largely attenuated and no PM recruitment is seen with *art1*^ΔPY^(Figure S3B, S3C) . We next tested whether Art1 facilitates PM recruitment of Rsp5. As expected, methionine treatment induces Rsp5 PM recruitment in cells expressing WT Art1, but this recruitment is much reduced in cells expressing *art1*^K486R^ (Figure S3E, S3F). Taken together, our results support the model that specific ubiquitination of adaptor proteins is required for proper recruitment of Rsp5 to target membranes and subsequent ubiquitin-mediated endocytosis of cargo proteins.

### Substrate dependent PM recruitment of adaptor protein Art5 and Art1

Next, we sought to examine if cargo proteins are required for adaptor protein recruitment to their functional locations. To do so, we examined the Art5-GFP PM recruitment in *ITR1-WT* and *itr1*Δ mutants upon inositol treatment for 1 hour. Strikingly, we found that the PM recruitment of Art5-GFP is abolished in the *itr1*Δ mutant (Figure 3G, 3H). Similarly, we observed that the PM recruitment of Art1 is attenuated in the *mup1*Δ mutant with methionine induction for 1 hour (figure S3G and S3H). We further tested whether Art1 can be recruited to the PM in cells expressing *mup1*-Q49R mutant upon methionine treatment. Previous data showed that Mup1 mutant Q49R is unable to be endocytosed with methionine treatment (Guiney *et al*., 2016) and the *mup1-*Q49R mutation abolishes the protein-protein interaction between Mup1 and Art1 (Figure S2E). We expected that PM recruitment of Art1 will be impaired in this mutant due to the loss of the interaction between Art1 and Mup1-Q49R. Indeed, PM recruitment of Art1 is abrogated in the *mup1*-Q49R condition (Figure S3G and S3H), suggesting that the Mup1-Art1 interaction is required for methionine-induced Art1 PM re-localization. Collectively, our data demonstrate that the substrate proteins (Itr1 and Mup1) are required for adaptor protein (Art5 and Art1) recruitment to target membranes (Fig. 3I).

### K63-linked di-ubiquitination enhances the interaction between adaptor proteins and Rsp5

We found that the Rsp5 adaptor proteins (Art5, Art1, and Art4) undergo K63-linked di-ubiquitination and this modification is required for efficient recruitment of Rsp5 to target membranes and cargo protein degradation. We hypothesized that adaptor di-ubiquitination enhances protein-protein interactions between di-ubiquitinated adaptors and Rsp5 and thus promotes the recruitment of the E3 ligase. To test this hypothesis, we set out to examine the binding between mono-Ub or K63-linked di-Ub and the HECT domain of Rsp5, as well as the binding between PY motifs and WW domains. To do so, we first generated K63-linked ubiquitin chains using K63-chain specific E2 enzymes Mms2/Ubc13 (Hofmann & Pickart, 1999; Sato *et al*, 2008; Spence *et al*, 1995). Then, we performed a binding assay between glutathione-*S*-transferase (GST) fusion proteins to Rsp5 HECT domain or GST only and the K63-linked ubiquitin chains. The mono-Ub and K63-linked di-ubiquitin chains bind to GST-HECT domain (lane 7), but not to GST (Figure 4A). The binding between mono-Ub and HECT domain depends on the exosite/ubiquitin interface (Y516 and F618) (French *et al*, 2009; Kim *et al*., 2011); we found that the binding between K63-linked di-Ub and HECT domain is disrupted by the exosite mutants Y516A, F618A, or the Y516A/F618A double mutant (lane 8, 9 and 10 of Figure 4A), suggesting that the K63-linked di-Ub also interacts with the HECT domain via the exosite.

**Figure 4.**
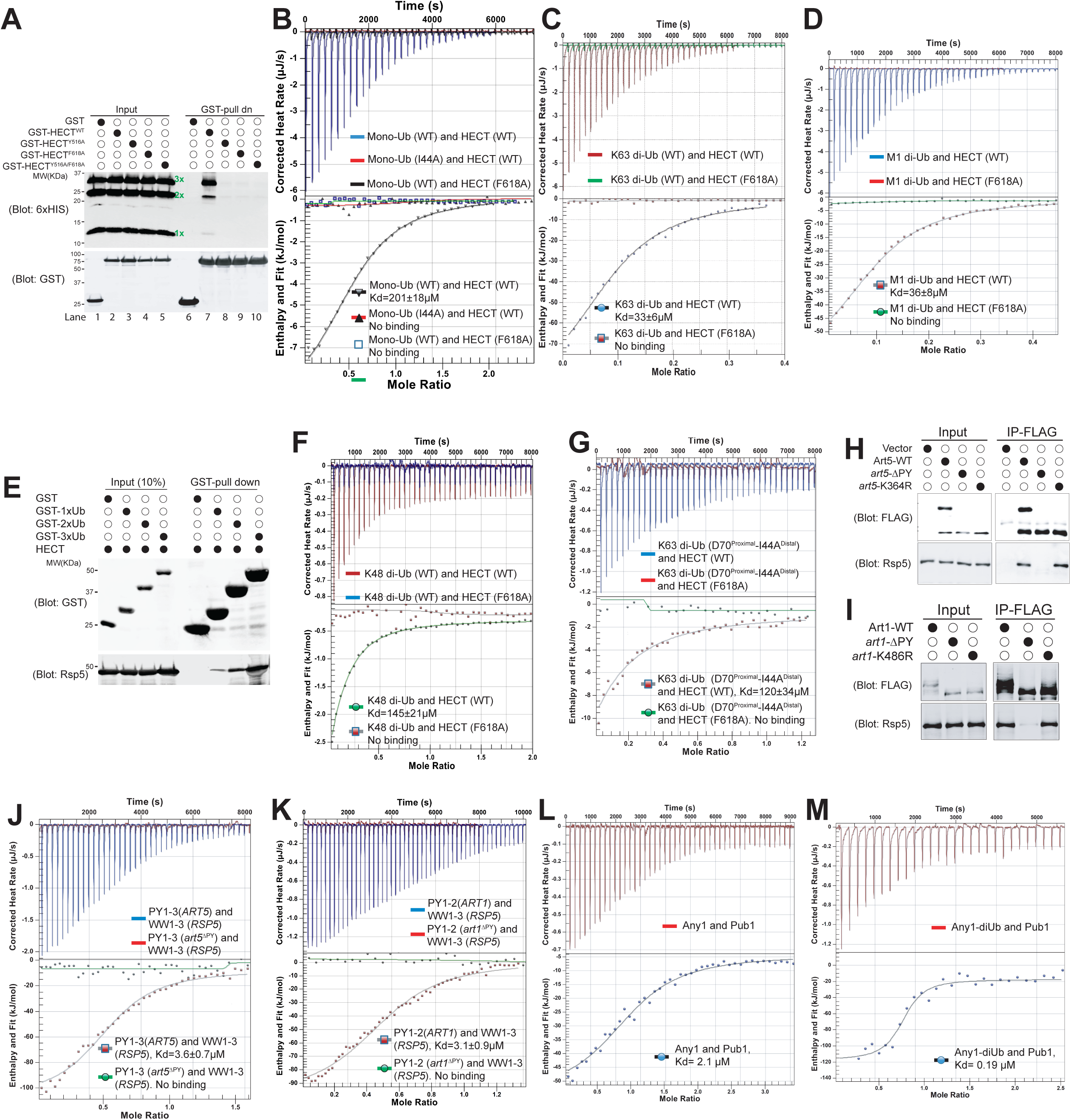
K63-linked di-ubiquitination enhances the interaction between adaptor proteins and Rsp5. (A) GST pull down assay between HECT-WT or F618A mutant and K63 linked Ubiquitin ladder. (B) Example ITC titration curves showing the binding of Mono-Ub-WT or I44A mutant to Rsp5 HECT domain. (C) ITC-based measurements of the bindings between K63 di-Ub and Rsp5 HECT domain. (D) The representative ITC curves of showing the binding of M1 linked di-Ub and Rsp5 HECT domain. (E) GST pull down assay between GST only, GST-1xUb, 2xUb or 3xUb and Rsp5 HECT domains. (F) Measurement of affinity between K48 di-Ub and Rsp5 HECT domain by ITC. (G) ITC-based measurements showing that the K63 di-Ub with a distal end ubiquitin mutant (I44A) partially disrupts the binding affinity with Rsp5 HECT domain. (H-I) IP of Art1 and Art5, WT, KR and PY-motif mutants with Rsp5-HECT domain. (J-K) ITC analysis of Art1 or Art5 PY motifs containing domain and Rsp5 WW1-HECT domain. (L) Analysis of binding affinity between Any1 (Art1 orthologue in Pombe) and the Pub1 (Rsp5 orthologue in Pombe). (M) ITC results obtained by titration of Any1 conjugated with K63 di-Ub into a solution of Pub1.

Each Need4 family E3 ligase contains a HECT domain. It was shown that HECT domains of various Nedd4 family HECT E3 ligases (Maspero *et al*., 2011), as well as the Rsp5 HECT domain (Kim *et al*., 2011), are able to interact with mono-Ubiquitin. Since we have shown that adaptor proteins are di-ubiquitinated in a K63-linkage, we next decided to examine the interaction between HECT domains and mono-Ub and K63-linked di-Ub. The dissociation constant (Kd) for the interaction between HECT and mono-ubiquitin was quantified by isothermal titration calorimetry (ITC) assay to be approximately 201µM (Figure 4B). We also employed the Rsp5 HECT exosite mutant (F618A) as a negative control. In agreement with the *in-vitro* GST-binding assay result in figure 4A, no binding was detected between mono-ubiquitin and HECT domain mutant (F618A, shown in the figure 4B). Ubiquitin is often recognized through a hydrophobic surface containing Ile44, which is bound by most Ubiquitin Binding Domains (UBDs) (Dikic *et al*, 2009; Shih *et al*, 2000; Sloper-Mould *et al*, 2001). We therefore included the ubiquitin binding mutant (I44A) serving as a negative control here. As expected, the I44A mutation of ubiquitin abolishes the binding between mono-ubiquitin and the HECT domain (Figure 4B). Our results suggest that the HECT domain exosite and the I44-containing ubiquitin hydrophobic surface are required to bridge the protein-protein interaction between the HECT domain and ubiquitin. In contrast to the mono-ubiquitin results, K63-linked di-Ub enhances the binding affinity Kd=33µM, nearly 6-fold relative to the mono-Ub (Figure 4C). In line with the *in-vitro* GST binding result, we examined the binding between K63-linked di-Ub and the HECT domain mutant (F618A) by ITC and found that the protein-protein interaction is abolished. Head-to-tail M1-linked di-Ub was proposed to mimic the K63 ubiquitin linkage (Komander *et al*, 2009; Zhu *et al*, 2017). As expected, our ITC analysis showed that M1 linked di-Ub binds to HECT with Kd=36µM, comparable with the K63-linked di-Ub (Fig. 4D). In line with this result, our in-vitro binding assay showed that the binding between GST-2xUb and HECT domain is stronger than GST-Ub (Fig. 4E). In comparison, K48-linked di-Ub shows a much lower affinity than K63-di-Ub, Kd=145µM (Fig. 4F). Together, our results demonstrate that HECT domain specifically binds to linear form K63 di-Ub and the exosite site is required for ubiquitin binding.

We next wondered if both the proximal and distal end ubiquitin of the K63-linked di-Ub, or just the distal end Ub, contribute the binding to the HECT domain. Since Ile44 of ubiquitin is essential for binding of ubiquitin to HECT domain (Fig.4B), we fused a distal end Ub (I44A) mutant to a proximal Ub (WT) and generated the distal end I44A mutant of K63 di-Ub (Ub^I44A^-Ub^WT^). The Ile44 residue of the proximal end ubiquitin is essential for ubiquitin binding by Ubc13/Mms2 and critical for K63-linked di-Ub catalysis, the Ile44 mutant of the proximal end ubiquitin of the K63 di-Ub cannot be made (Tsui *et al*, 2005). We found that the K63-linked Ub^I44A^-Ub^WT^ binds to HECT with an Kd=120µM, lower binding affinity than the K63 di-Ub (Fig. 4G). Thus, our result suggests that both distal and proximal ubiquitins contribute to the HECT domain binding, probably cooperatively.

We next sought to determine if K63-linked di-Ub enhances the binding between adaptor and HECT type E3 ligase. In spite of the fact that KR mutants of Art5 and Art1 lead to attenuated Rsp5 PM recruitment and cargo proteins (Itr1 and Mup1), we still observed the interaction between Art1-K486R or Art5-K364R with Rsp5 using Co-IP (Fig. 4H, 4I), probably due to the interaction between the PY motifs and WW domains. Indeed, as shown in the figure (Fig. 4J, 4K), Art1 and Art5 PY motif containing peptides interact with purified WW domains from Rsp5 (Kd= 3.6µM for Art1 PY motifs and Kd=3.1µM for Art5 PY motifs), but not with PY motif mutants. Since the sole interaction between PY motifs and WW domains does not suffice the full activation of Rsp5 function (Fig. 2A and S2A), the interaction between di-Ub and HECT may enhance the binding affinity between Need4/Rsp5 E3 ligases and their adaptors. We next sought to test the binding between full length adaptors and Rsp5. We found that we could not express Art1 or Art5 at high levels in *E. coli,* then tried to express the Art1 orthologue from *S. pombe,* Any1, in *E.coli*. We found the *S. pombe* Rsp5 ortholog Pub1 interacts with Any1 with a binding affinity Kd∼2.1µM (Fig. 4L), in a similar range as the binding affinity between PY motifs and WW domains shown earlier (Fig. 4J, 4K). Remarkably, Any1 conjugated with K63 di-Ub enhances the binding with Pub1 over 10-fold in comparison with non-conjugated Any1 (Figure 4M), suggesting that di-Ub conjugation onto Any1 probably leads to a structural conformation change of Any1 and therefore enhances the binding with Pub1. This result is in agreement with our previous result that di-ubiquitination of Art5 and Art1 are required for efficient Rsp5 recruitment to the plasma membrane and for cargo protein sorting. Taken together, the di-ubiquitination of adaptor proteins enhances the binding affinity with the E3 ligase, leading to E3 ligase recruitment and cargo protein ubiquitination and sorting.

### Deubiquitination of K63 di-Ub of adaptor protein Art5 by Ubp2

Given that the exosite of Rsp5 is essential for binding with the di-Ub on adaptor proteins, we next examined the ubiquitination status for the adaptor proteins Art1 and Art5. The di-ubiquitinated form of Art5 is diminished in the *rsp5*-F618A mutant (Fig. 5A, lane 2). Similarly, the di-ubiquitinated pool of Art1 is substantially attenuated in either the Y516A or F618A exosite mutant (Fig. S4A). Maspero and coworkers reported that exosite mutants do not alter the binding affinity between E3 and E2 enzymes, the transthiolation process from E2 to E3, or the self-ubiquitination activity of Nedd4 (Maspero *et al*., 2011). We therefore speculated that a deubiquitination enzyme (DUB) is involved in the trimming process of the K63-linked di-Ub. To test this hypothesis, we performed a multicopy gene suppression screen with all budding yeast DUBs . As shown in Figure S4B, overexpressing Ubp2 by a *TDH3* promoter leads to a reduction of the di-ubiquitinated portion of Art1. Further, we overexpressed the catalytic dead mutant C745V of Ubp2 and found that the deubiquitination of Art1 is restored (Fig. S4C). This result infers that Ubp2 may function as a DUB to trim the di-ubiquitinated form of Rsp5 adaptor proteins.

**Figure 5.**
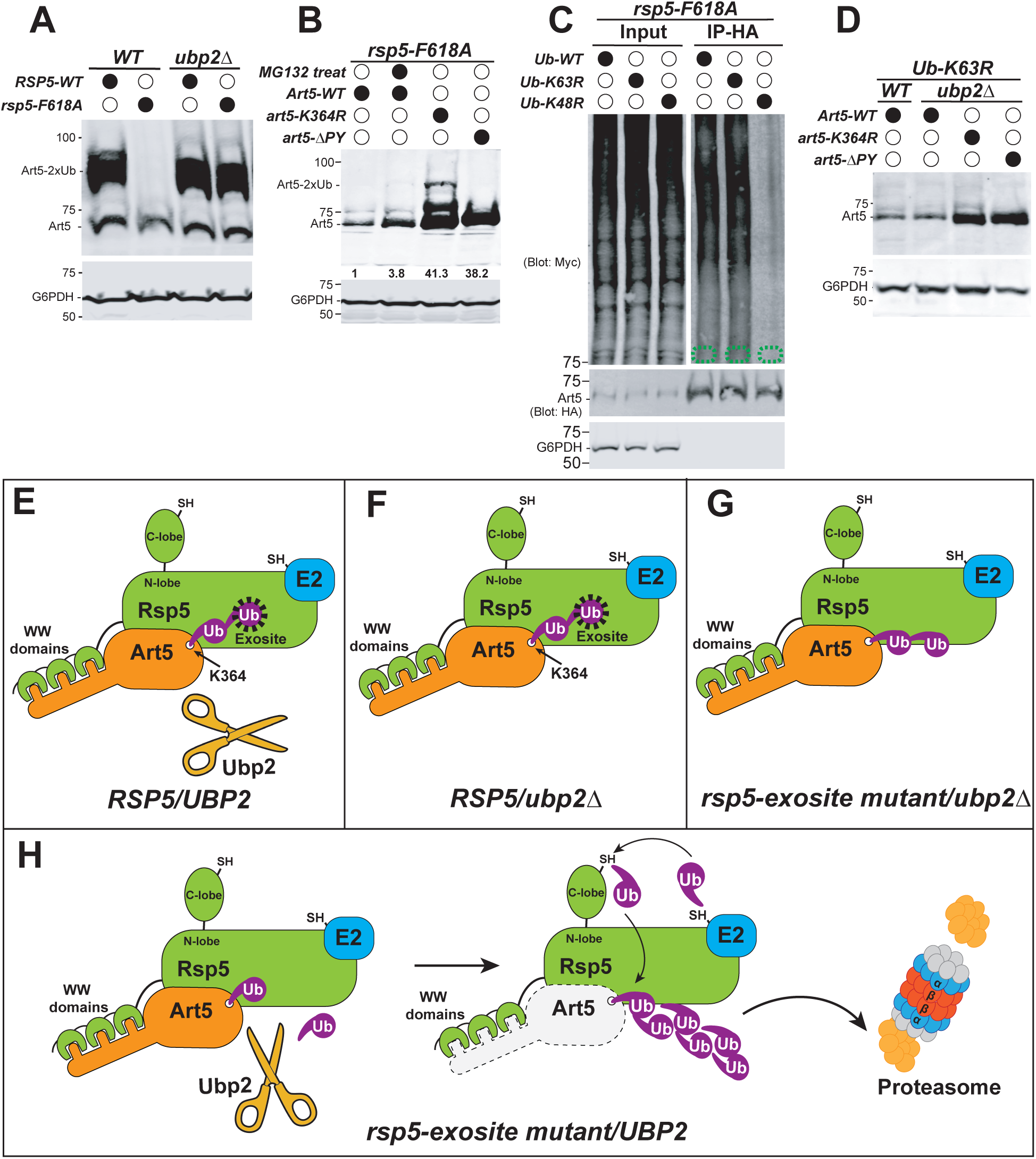
Deubiquitination of Art5 di-Ub by Ubp2. (A) Immunoblot analysis of Art5-3HA in the indicated yeast strains: RSP5(WT*), rsp5-*F618A, ubp2Δ and *rsp5-F618A*/ubp2Δ. (B) Yeast strain *rsp5-*F618A expressing Art5-3HA, *art5*^K364R^-3HA and *art5*^ΔPY^-3HA were mock treated with DMSO and the cells bearing Art5-3HA were treated with MG132 (25µg/ml) for 60min. (C) Ubiquitin blot of *rsp5-*F618A yeast cells carrying Art5-3HA, as well as WT, K63R or K48R myc-ubiquitin expression vector. Cells were treated with MG132 (25µg/ml) for 60 min. Samples were immunoprecipirated using anti-HA antibody and analyzed by immunoblot. (D) Yeast mutants *Ub-K63R* and *Ub-K63R*/*ubp2Δ* expressing Art5-3HA, *art5*^K364R^-3HA and *art5*^ΔPY^-3HA. (E-H) models depicting that Ubp2 and Rsp5 modulates the K63 di-Ub and K48 polyubiquitination of Art5 together: (E) K63 di-Ub of Art5 is protected from Ubp2 cleavage when engaged into exosite in *RSP5/UBP2* condition. (F) Art5 remains engaged in exosite as K63 di-Ub in *ubp2*Δ mutant. (G) Art5 is not engaged in the exosite but kept as K63 di-Ub in *rsp5-F618A*/ubp2Δ condition. (H) K63 di-Ub of Art5 is cleaved by Ubp2 and K48 polyubiquitin chain is instead conjugated at the K364 of Art5 by Rsp5 before proteasomal degradation.

To investigate the role of Ubp2 in the modification of Rsp5 adaptor proteins, we examined the adaptor protein Art5 in a double mutant of *rsp5*-exosite (F618A) and *ubp2*Δ. Strikingly, the di-ubiquitinated Art5 and Art1 are nearly fully restored in the *rsp5*-F618A/*ubp2*Δ strain (Fig. 5A,lane 4) and (Fig. S4D, lane 4), indicating that Ubp2 trims the di-ubiquitin on adaptors Art5 and Art1 when they are disengaged from the Rsp5 exosite. To test if Ubp2 is playing a catalytic or structural role in this process, we complemented the *rsp5*-F618A/*ubp2*Δ with either a wild-type or a catalytic mutant *ubp2*-C745V. We found that the Ubp2-WT (Fig. S4E lane 2) fully reverses the rescue of Art1 trimming seen in the lane 1, whereas the *ubp2*-C745V does not (Fig. S4E lane 3). Together, these results suggest that the exosite can protect the di-ubiquitin moiety on adaptors from the cleavage by Ubp2.

We next wondered if the loss of Art5 (Fig. 5A, lane 2) is mediated by proteasome function. To answer this question, we treated the *rsp5*-F618A mutant with proteasome inhibitor MG132. We found that the full length Art5 protein is restored 2.8 fold with temporary inhibition of proteasome function (Fig. 5B), suggesting that Art5 probably undergoes K48-linked polyubiquitination because K48-linked ubiquitin chains are is preferred by the proteasome. As seen in figure 5B, either the PY motif or the K364R mutant rescues the loss of Art5, indicating that Rsp5 is the E3 ligase responsible for Art5 degradation and the same site K364 is used for this ubiquitination process. To directly determine the involvement of K63 versus K48 linkage in the Art5 degradation, we examined the effect of overexpressing myc-ubiquitin with wild-type, K63R and K48R mutations on Art5 ubiquitination. We found that expressing myc-ubiquitin K63R does not affect Art5 hyperubiquitination in the *rsp5*-exosite mutant background, whereas the K48R ubiquitin mutant substantially reduced Art5 ubiquitination (Fig. 5C). This result suggests that the Art5 ubiquitination in the *rsp5*-exosite mutant is mediated by a K48-linked polyubiquitin chain.

We then sought to uncover the mechanism by which the Art5 degradation is triggered. We observed that Art5 protein is also degraded in the *Ub*-K63R mutant (Fig. 1C). We wondered if *ubp2*Δ can rescue Art5 degradation in the *Ub*-K63R mutant. To test this, we deleted Ubp2 in the *Ub*-K63R mutant and found that *ubp2*Δ does not reverse the loss of Art5 protein (Fig. 5D). Our results suggest that the loss of K63-di-Ub on Art5, instead of Ubp2, in either the *Ub*-K63R mutant or *rsp5*-exosite mutant, leads to Art5 degradation. Remarkably, in these two conditions, the Art5 degradation can be rescued in *art5*-K364R and *art5*-ΔPY mutant (Fig. 5B and 5D). These data suggest that Rsp5 can mediate both K63 and K48-linked ubiquitination and the K364 residue of Art5 can be conjugated with both K63-linked di-Ub and K48-linked polyubiquitin chain. Collectively, our results support a working model that the K63-linked di-Ub on Art5 is fully engaged into the Rsp5 exosite so that Ubp2 cannot cleave it efficiently, whereas the K63-linked di-Ub is disengaged in an *rsp5*-exosite mutant therefore exposed to Ubp2 for cleavage. Upon the cleavage of K63-linked di-Ub, a K48-linked polyubiquitin chain is conjugated at the same residue Art5-K364 thereby leading to the proteasome-dependent degradation of Art5 (Fig. 5E-5H).

Since PM recruitment of Rsp5 is enhanced by Art5 or Art1 protein ubiquitination (Fig. 3E, S3E) and Ubp2 can deubiquitinate these adaptor proteins (Fig. 5A and S4D), we therefore asked if the adaptor ubiquitination process is reversible and Ubp2 is involved in this process or not. To monitor the pre-existing adaptor proteins, we decided to employ the *tet*-Off system to fix the pool of adaptor proteins by treating the cells with doxycycline. As seen in the figure S4F and S4G, pre-existing Art5 or Art1 undergoes ubiquitination upon inositol or methionine treatment for 1 hour in both WT and *ubp2*Δ conditions, whereas the adaptor proteins shifted back to less ubiquitinated status after removing the stimulation in the WT condition, but not in the *ubp2*Δ mutant. Together, our data demonstrate a model for adaptor protein recycling mediated by Ubp2 (Fig. S4H). First, stimulation enhances adaptor ubiquitination. Second, the ubiquitinated pool of adaptor proteins can be de-ubiquitinated by Ubp2 when stimulation is terminated.

## Discussion

In this study, we identified the first K63-linked di-Ub modification that modulates the function of Rsp5 and adaptor proteins. Our data demonstrates that two biological functions are implicated with this K63-linked di-Ub modification. First, K63-linked di-Ub activates Rsp5 function. K63-linked di-Ub enables the full engagement of adaptors onto the Rsp5 exosite and sharply enhances the binding affinity with Rsp5, which facilitates Rsp5 recruitment and accelerates substrate protein ubiquitination. Second, K63-linked di-Ub prevents the adaptors from being conjugated with K48-linked polyubiquitin. K63-linked di-Ub on adaptors engaged with the Rsp5 exosite are not accessible to Ubp2. Once released from Rsp5 exosite, the exposed K63-linked di-Ub is subjected to cleavage by Ubp2 and K48-linked polyubiquitin subsequently can be conjugated onto the adaptor protein, which signals proteasome-dependent protein degradation. Further, we monitored the ubiquitination status of adaptor proteins Art1 and Art5. Using *tet*-Off system, we have shown that adaptor proteins undergo ubiquitination upon stimulation and Ubp2 is required for deubiquitination of adaptor proteins once the stimulation is removed. As hypothesized by our earlier review (MacGurn *et al*, 2012), our current data supports the model that ubiquitinated adaptor proteins were deubiquitinated by Ubp2 once released from Rsp5 exosite so that the adaptor proteins can be recycled for the next round of ubiquitination event.

### K63-linked di-Ub is engaged into Rsp5 E3 ligase for activation

While we showed that Rsp5 adaptors Art1, Art4 and Art5 undergo K63-linked di-Ub modification, we also demonstrate that this conjugation sharply enhances the binding with the E3 ligase and activates the E3 ligase function for substrate ubiquitination (Fig. 2D). We reason that the interaction between the di-Ub chain and the HECT domain locks the E3 ligase and adaptor into an active/functional conformation. For adaptor-independent ubiquitination, the Nedd4/Rsp5 ligase exosite is also required for efficient ubiquitin conjugation, demonstrating that the “Ub-exosite binding” is required to localize and orient the distal end ubiquitin chain to promote conjugation (Kim *et al*., 2011; Maspero *et al*., 2011). In terms of the Rsp5 adaptor-mediated function, we propose that the binding between “di-Ub and exosite” not only enhances the binding affinity between the E3 ligase and adaptor (Fig. 4L-4M), but also leads to more productive Rsp5 recruitment to properly orient and present the substrate for ubiquitination at target membranes (Fig. 3B).

While we presented the evidence of E3 ligase activation by ubiquitinated adaptors, we also showed that K63 di-Ub generates a 6-fold tighter binding to the HECT domain than mono-Ub. We reason that the K63 di-Ub provides alternative options to bind a single site, but also fits with a model in which there are multiple ubiquitin binding sites. It was found that three N-lobe mutations (Y516A, F618A, and V621A/V622A) completely abolished ubiquitin binding and three extra mutations (N513A, Y521A, and R651A) caused a reduction in binding (French *et al*., 2009). Kim and coworkers found that the L8-I44-V70 hydrophobic patch of mono-Ub sits on Rsp5 in three legs, like a tripod (Kim *et al*., 2011). Likewise, two separated UIMs in Rap80 bind to extended K63-linked ubiquitin chain favorably (Sato *et al*, 2009; Sims & Cohen, 2009). Indeed, we have shown the results that K63-linked di-Ub with a mutation (I44A) at the distal end ubiquitin leads to lower binding with Rsp5 (Figure 4G). We propose that multiple ubiquitin binding sites are probably present at the Rsp5 exosite to accommodate the two hydrophobic patches of the distal and proximal ubiquitins, which needs be addressed in the future by structural analysis.

### The linkage specificity and length control for the K63-linked di-Ub

We have been intrigued by the question of how the K63 linkage of di-Ub was achieved and preferred, instead of K48. It is known that yeast Rsp5 and human Nedd4 mainly assemble K63-linked ubiquitin chains (Kim & Huibregtse, 2009; Maspero *et al*., 2011). The K48-linked di-Ub binds the HECT domain, but not as tight as K63-linked di-Ub (Fig. 4F). Interestingly, both the M1-linked and K63-linked di-Ubiquitins adopt an equivalent open conformation (Komander *et al*., 2009) and exhibit similar binding affinity to the HECT domain (Fig. 4D), indicating that the HECT domain exosite has a strong preference for the linear and extended form of di-Ub. In contrast, the K48-linked polyubiquitin chain adopts a significantly distinct and compact structure (Eddins *et al*, 2007), which may not be favorable for the HECT domain exosite.

Why is the K63-linked di-Ub chain limited to a dimer? On the one hand, this probably correlates with the physiological reversible function of adaptors. The K63-linked 3x or longer ubiquitin chains likely generate stronger binding with the HECT domain than di-Ub (Fig. 4E). We reason that the di-Ub binds well with the HECT domain, but still can be disengaged from the HECT domain under physiological conditions so that Rsp5 can be disassociated and recycled. On the other hand, the K63-linked di-Ub is probably just enough to be masked by the HECT domain exosite cavity whereas longer chains will be trimmed by Ubp2. Future structural studies could address the accessible region for the di-Ub isopeptide bond cleavage by Ubp2 when di-Ub is engaged into the HECT domain. Further, a K63-linked polyubiquitin chain also can serve as a targeting signal for proteasomal degradation (Ohtake *et al*, 2018; Saeki *et al*., 2009). We noticed that hyperubiquitinated forms (>2xUb) of ART proteins are not stables since the linear form of 3xUb leads to adaptor protein degradation (Lanes #7 of the figures 1C, S2C and S2F). Indeed, K63-linked polyubiquitin on Rsp5 adaptor proteins contributes to proteasomal degradation of the adaptors (Ho *et al*, 2017).

### Ubp2 mediates the recycling of Rsp5 E3 ligases from adaptors after ubiquitination

McDonald and coworkers proposed that several Rsp5 adaptors compete for Rsp5 and a Ubp2 deficiency increased both the adaptor activity and the ability to compete for Rsp5 (MacDonald *et al*., 2020). Indeed, the PPxY motif containing Rsp5 adaptors share the E3 ligase Rsp5 and an adaptor should disassociate from Rsp5 to allow other adaptors to engage with Rsp5 to ubiquitinate different substrate proteins. In agreement with this working model, Nedd4-mediated downregulation of the sodium channel ENaC is impaired when Nedd4 is sequestered by overexpression of another Nedd4 E3 adaptor, Ndfip2 (Konstas *et al*, 2002).

Besides cleavage of K63 di-Ub in the *rsp5*-exosite mutant, Ubp2 allows the recycling of Rsp5 from its adaptor proteins. Since K63 di-Ub greatly enhances the binding affinity between adaptors and E3 ligase (shown in figure 4), Ubp2 likely helps the Nedd4/Rsp5 E3 ligase to catalyze distinct ubiquitination events by cleaving the di-Ub off the adaptors and recycling Rsp5. The multitasking of Rsp5 via various adaptors leads us to hypothesize that activated Rsp5 can be released from engaged adaptor proteins. We showed that the adaptor proteins Art1 and Art5 undergo di-Ubiquitination upon environmental stimulation and Ubp2 is required to reverse this ubiquitination. Once the ubiquitination is done, the engaged K63 di-Ub is exposed for cleavage by Ubp2. Thereafter, Ubp2 acts on ubiquitinated adaptor proteins to release the adaptor proteins and Rsp5. The mechanism by which Ubp2 executes this reaction needs to be addressed in the future.

In summary, we propose that Rsp5 ubiquitinates adaptors to trigger their engagement with the Rsp5 exosite, which enables the tight binding between adaptors and Rsp5 thereby activating Rsp5 function. Ubp2 acts as an antagonist for K63 di-Ub to modulate the interaction between K63-di-Ub and the Rsp5 exosite in a reversible manner to maintain cellular homeostasis of Rsp5. Future work needs to address the atomic structure of the ART family of adaptor proteins in complex with Rsp5 in order to understand how di-Ub is attached to the adaptor and how the di-ubiquitinated adaptors engage with the HECT E3 ligases, stabilizing an activated conformation of the E3 ligase.

## Material and Methods

### Yeast strains, cloning, mutagenesis and cell growth conditions

The *ART1, ART4, ART5, ITR1, MUP1* and *YUH1* genes were cloned from *Saccharomyces cerevisiae* yeast strain SEY6210. Pub1 (residue 287-767) and Any1 (residue 17-361) were PCR amplified from *Schizosaccharomyces pombe* yeast strain PR109 and subcloned into pET28a with an N-terminal 6xHis-SUMO tag. When necessary, the gene deletions and taggings were made using gene replacement technique with longtine-based PCR cassettes (Longtine et al., 1998). All yeast strains and plasmids are described in Tables S1 and S2. For fluorescent microscopy experiments, cells were grown overnight to mid-log phase (OD600∼0.5) in synthetic media at 30°C. For inositol or methionine stimulation experiments, cells were grown in synthetic media to log phase (OD600∼0.8) then treated with exogenous inositol and methionine at different concentrations. Ub-WT, Ub-K63R, Ub-K48R, Ub-D77, Mms2 and Ubc13 were PCR amplified from yeast strain SEY6210 genomic DNA and cloned into pET21a, pET28a-6xHIS and pGEX6p-1 respectively. 1x, 2x, and 3x and 4x Ub head-to-tail fusions of Art1, Art4, Art5 expression and pGEX6p-1 vectors were made by Gibson assembly. E1 enzyme expression vector pET21a-Uba1 (human) and K48 ubiquitin linkage specific E2 enzyme E2-25K expression vector pGEX-6p-1-E2-25K are from our lab stock. YUH1 was subcloned into pGEX6p-1 expression vector with an N-terminal GST tag. PY motifs containing regions for Art1 (661-710) and Art5 (520-586) were PCR amplified and cloned into pGEX-6p-1 vectors. Rsp5 HECT domain (444-809) and WW1-HECT domain (224-809) were fused with N-terminal SUMO tag and cloned into pET28a vector.

### Protein Purification

All pET21a, pET28a, pGEX6p-1 constructs were transformed into *Escherichia coli* strain Rosetta (DE3) cells. Single colonies were then cultured in Luria-Bertani (LB) medium containing either 100µg/ml Ampicillin or 50 µg/ml kanamycin to a density between 0.6 and 0.8 OD600 at 37°C. Cultures were induced with 0.2 mM isopropyl-B-D-thiogalactopyranoside (IPTG) at 18°C for 16 hours. *E.coli* cells were collected by centrifugation at 3,500 rpm for 15 min at 4°C. For non-tagged ubiquitin purification, cells were disrupted by sonication in the lysis buffer (50mM NH4Ac (pH4.5rt), 2mM DTT, 1mM EDTA, 1mM PMSF). For 6xHIS-SUMO tagged proteins, cells were sonicated in the lysis buffer (20 mM Tris (pH 7.5), 150 mM NaCl, 2mM DTT, 1mM EDTA, 1mM PMSF). For GST fusion proteins, cells were disrupted in the lysis buffer (200mM NaCl, 25mM Tris.HCl pH8rt, 2mM EDTA, 2mM DTT, 1mM PMSF).

The lysate for Ub (WT, K63R, K48R, I44A, D77 or D77/I44A) was adjusted to pH4.5 then spun down at 46,000xg for 45 min at 4°C. The supernatant was heated at 70°C for 5 min then spun down again with the same condition. The supernatant was loaded onto SP Sepharose Fast Flow resin pre-equilibrated with the same lysis buffer (pH4.5). The Ub was eluted with 50mM ammonium acetate (pH4.5 room temperature) buffer containing 2mM DTT using a linear gradient of 0-500mM NaCl. The eluted Ub mutants were fractionated by Superdex 200-exclusion column then dialyzed against size-exclusion buffer (20 mM Tris (pH 7.5), 150 mM NaCl, 2mM DTT). Each mutant was concentrated to 15mg/ml and stored at -80 °C.

For 6xHIS-SUMO-tagged (HECT, Pub1(287-767), Any1(17-361) and WW1-HECT) and GST-tagged proteins (Ubc13, E2-25K, Yuh1, PY motifs of Art1 or Art5 and M1 linked Ub-Ub), the sonicated lysates were centrifuged 46,000xg for 45 min at 4°C. The supernatant was bound with TALON cobalt resin or Glutathione Sepharose 4 Fast Flow and the resins were digested by SUMO-specific Ulp1 or GST-specific PreScission proteases to release the proteins of interest. The eluted proteins were fractionated by Superdex 200 using size-exclusion buffer (20 mM Tris (pH 7.5), 150 mM NaCl, 2mM DTT). Ubc13, E2-25K and Yuh1 were concentrated to 750µM with 20% glycerol and the other proteins were concentrated to 1mM and stored at -80°C.

For 6xHis-tagged Uba1 and Mms2 purification, the *E.coli* cells were sonicated in lysis buffer 20 mM Tris (pH 7.5), 150 mM NaCl, 2mM DTT, *c*Omplete^TM^ protease inhibitor). The cell lysate (per 1 liter) was cleared by centrifugation at 46,000xg, 45min, 4°C. The supernatant was incubated with cobalt-chelate TALON resin for 30min before column wash with lysis buffer supplemented with 25mM imidazole and the protein of interest was eluted with 300mM imidazole and dialyzed against 50mM Tris-HCl (pH7.6) containing 2mM DTT and 0.1mM EDTA. The protein is concentrated to 100µM with 20% of glycerol and stored at -80°C.

For GST-tagged protein (GST-1xUb, GST-2xUb and GST-3xUb) purification, the sonicated cell lysate was spun down at 46,000xg, 45min, 4°C. The supernatant per 1 liter of cells was incubate with 2ml of Glutathione Sepharose 4 Fast Flow resin and washed with 5 column volumes of wash buffer (20mM Tris pH8rt, 200mM NaCl, 1mM DTT). The GST-tagged proteins were eluted by 2 column volumes of elution buffer (100mM Tris pH8.5, 20mM Glutathione) then dialyzed against size-exclusion buffer (20 mM Tris (pH 7.5), 150 mM NaCl, 2mM DTT). Each protein was concentrated to 30mg/ml and stored at -80oC.

For synthesis of K63 or K48 di-Ub proteins, 5xPBDM buffer was prepared: 250 mM Tris-HCl (50%, pH 8.0, or pH7.6), 25 mM MgCl_2_, 50 mM creatine phosphate (Sigma P7396), 3 U/mL of inorganic pyrophosphatase (Sigma I1891), and 3 U/mL of creatine phosphokinase (Sigma C3755). K63 linked di-Ub is synthesized by incubating purified human E1 (0.1µM), yeast E2 (Ubc13 and Mms2, 8µM of each), two ubiquitin mutants (K63R and D77, 5mg/ml of each), ATP (2.5mM), 1 mM DTT and 1xPBDM buffer (pH7.6). For K48 linked di-Ub synthesis, purified human E1 (0.1µM), E2-25K (20µM), two ubiquitin mutants (K48R and D77, 7.5mg/ml of each), ATP (2.5mM), 1 mM DTT and 1xPBDM buffer (pH8.0) were mixed. The reaction mixtures of either K63 or K48 di-Ub were incubated at 37°C for overnight then the reaction was chilled on ice for 10min to stop the reaction. 0.2 volume of 2M ammonium acetate was added to the reaction to decrease the pH to less than 4.0. The mixture were loaded to SP Sepharose Fast Flow. The K63 di-Ub or K48 di-Ub mixtures were loaded onto Superdex 75 size-exclusion column using gel filtration buffer (20mM Tris-HCl (pH7.5), 2mM DTT, 150mM NaCl) and the fractions of diUb were pooled and concentrated.

### Synthesis and Purification of Any1-diUb

To remove the D77 of the proximal Ub and unlock the carboxyl-terminal Gly-Gly of K63diUb for further conjugation, purified K63 linked di-Ub (30mg/ml) is exchanged into hydrolysis buffer (50 mM Tris-HCl pH 7.6, 1 mM EDTA, and 1mM DTT) and treated with purified Yuh1 (final concentration of 16µg/ml) for 60minutes at 37°C. After cooling down the reaction at room temperature, 4mM DTT to the mixture is supplememted with DTT to 5mM (final concentration). The reaction mixture was then applied to a 5 ml Q column equilibrated with Q buffer (50 mM Tris-HCl pH 7.6, 1 mM EDTA, 5 mM DTT). After 2 bed volumes of wash, the unbound K63 di-Ub (D77 removed) is collected and concentrated. Di-ubiquitination of Any1 was carried out by incubating purified Any1 proteins with human E1(0.1 µM), human E2(UbcH5C, 0.3 µM) and Pub1 (0.3 µM), K63 diUb (D77 removed, 10 µM), ATP (2.5mM), 1 mM DTT and 1xPBDM buffer (pH7.6) for 30min at room temperature. The reaction mixture was chilled on ice before loading onto Superdex 200 size-exclusion column using gel filtration buffer (150 mM NaCl, 20 mM HEPES pH 7.5), and fractions of Any1-diUb were pooled and concentrated.

### GST pull down assay

For pull-down experiments, 2 µM of GST fusion proteins were immobilized onto 100µL of glutathione bead slurry in the 1ml of pull down buffer (50mM Na-HEPES pH7.5, 150mM NaCl, 1mM EDTA, 1mM EGTA, 10% Glycerol, 1% Triton X-100). 500ng of Rsp5 HECT protein was added to the mixture and incubated at 4°C for 2 hours. After 4 washes with pull down buffer, specifically bound proteins were eluted by SDS-sample buffer and resolved on SDS-PAGE (11%) and detection was obtained by Coomassie-staining.

### Isothermal Titration Calorimetry assay

Isothermal Titration Calorimetry (ITC) experiments were carried out on an Affinity-ITC calorimeter (TA instruments) at 25°C. Titration buffer contained 20 mM Tris-HCl (pH 7.5), 150 mM NaCl, 1 mM DTT. For a typical experiment, each titration point was performed by injecting a 2 μL aliquot of protein sample (50–1000 μM) into the cell containing 300µL of another reactant (5–300 μM) at a time interval of 200 s to ensure that the titration peak returned to the baseline. The titration data was analyzed with NanoAnalyze v3.12.0 (TA instruments) using an independent binding model.

### Fluorescence microscopy assay

For fluorescence microscopy, cells expressing GFP, pHluorin or mCherry proteins were visualized using a DeltaVision Elite system (GE), equipped with a Photometrics CoolSnap HQ2/sCMOS Camera, a 100×objective, and a DeltaVision Elite Standard Filter Set (‘FITC’ for GFP/pHluorin fusion protein and ‘mCherry’ for mCherry fusion proteins). Image acquisition and deconvolution were performed using Softworx.

### Whole cell lysate extraction and western blotting

Whole cell extracts were prepared by incubating 6 ODs of cells in 10% Trichloroacetic acid on ice for 1 hour. Extracts were fully resuspended with ice-cold acetone twice by sonication, then vacuum-dried. Dry pellets were mechanically lysed (3x 5min) with 100 µL glass beads and 100 µL Urea-Cracking buffer (50 mM Tris.HCl pH 7.5, 8 M urea, 2% SDS, 1 mM EDTA). 100μl protein 2x sample buffer (150 mM Tris.HCl pH 6.8, 7 M urea, 10% SDS, 24% glycerol, bromophenol blue) supplemented with 10% 2-mercaptoethanol was added and samples were vortexed for 5 min. The protein samples were resolved on SDS-PAGE gels and then transferred to nitrocellulose blotting membranes (GE Healthcare Life Sciences).

The flowing antibodies and dilutions were used in this study: Rabbit polyclonal anti-G6PDH (1:30,000; SAB2100871; Sigma), Rabbit polyclonal anti-GFP (1:10,000; TP401; Torrypines), Mouse monoclonal anti-GFP (1:1,000; B-2, sc-9996; Santa Cruz), Mouse monoclonal anti-Myc (1:5,000, sc-40, Santa Cruz), IRDye® 800CW Goat anti-Mouse (1:10,000; 926-32210; LI-COR), IRDye® 800CW Goat anti-Rabbit (1:10,000; 926-32211; LI-COR), IRDye® 680LT Goat anti-Rabbit (1:10,000; 926-68021; LI-COR) and IRDye® 680LT Goat anti-Mouse(1:10,000; 925-68070; LI-COR).

### Immunoprecipitation (IP) assay

100 ODs of cells were collected and washed with water at 4°C. To examine the interaction between Art1 and Mup1-GFP, between Art5 and Itr1-GFP, or between ARTs protein and Rsp5. Yeast cells were washed with ice-cold water 3 times. The cells were lysed in 500 µl of IP buffer (20 mM Tris.HCl, pH 7.5, 0.5 mM EDTA, pH 8.0, 0.5 mM EGTA, 0.5 mM NaF, 150 mM NaCl, 10% glycerol, 1 mM PMSF, 10 mM *N*-ethylmaleimide (NEM), and *c*Omplete Protease Inhibitor). Cell extracts were prepared by glass-bead beating with 0.5-mm zirconia beads for five cycles of 30 seconds vortexing with 1 minute breaks on ice. Membrane proteins were solubilized by adding 500 µl of 1% Triton X-100 in IP buffer. The lysates were incubated at 4°C for 30 min with rotation then spun at 500xg for 5 min at 4°C. The supernatant was clarified by centrifugation at 16000xg for 10 min. To detect the interaction between ARTs and Mup1 or Itr1-GFP proteins, the cleared lysate was incubated with 50µl of GFP-nanotrap resin for 2 hours at 4°C. To examine the interaction between Rsp5 and ARTs, the cleared lysate was bound with 50µl of FLAG-M2 resin (Sigma, A2220) at 4°C for 2 hour. After incubation, the resin was washed 5 times with 0.1% Triton X-100 in IP buffer and the bound protein was eluted by 50 µl of 2x sample buffer.

To examine the ubiquitination of Itr1, Cells were grown to early log phase in synthetic media. Yeast strain (*doa4Δpep4Δart5Δ,* Itr1-GFP) cells co-expressing Myc-Ub expression vector (Zhu et al., 2017) and Art5^WT^ or *art5^K364R^* were grown to mid-log phase in synthetic medium at 30°C. Cells were pretreated with 0.1µM CuCl2 for 4 hours to induce the Myc-Ub expression prior to inositol (20µg/ml) treatment. 100ODs of Cells were incubated with 10% TCA buffer and the extracts were washed with cold acetone. Dry pellets were mechanically lysed (3x 5min) with 100 µL glass beads and 100 µL Urea-Cracking buffer (50 mM Tris.HCl pH 7.5, 8 M urea, 2% SDS, 1 mM EDTA, 200mM NEM). The cell lysates were mixed with 1ml of IP buffer (50 mM HEPES-KOH, pH 6.8, 150 mM KOAc, 2mM MgOAc, 1mM CaCl_2_, 20mM NEM and 15% glycerol) with cOmplete™ protease inhibitor (Sigma-Aldrich, St. Louis, MO). The Cell lysates were clarified by spinning at 16,000x*g* for 10 min at 4°C. The resulting lysate was then incubated with 50 µL GFP-nanotrap resin for 4 hours at 4°C. The resin was washed 5 times with 0.1% Triton X-100 in IP buffer. Bound protein was eluted by 50 µl of 2x sample buffer. Whole cell lysate and the IP reaction was resolved on 10% SDS-PAGE gels and the blots were probed with both GFP and Myc antibodies.

### Quantification of westernblot band intensity

Westernblot in figures were quantified using Image-J software. The significance for protein densities were determined two-tail *t*-test, α=0.05 (Bonferroni correction), n=3. n.s. indicates not significant; *, *P* < 0.05; **, *P* < 0.01; ***, *P* < 0.001.

### Quantification of microscopy images

Images of GFP-Rsp5, Art5-GFP and Art1-mNG were taken by fluorescence microscopy. The fluorescence signal of the target proteins at PM were selected and measured by Image-J. The corrected total fluorescence of each selection = Selected density **—** (Selected area X Mean fluorescence of background readings). The ratio of GFP-Rsp5, Art5-GFP and Art1-mNG recruitment to PM or vacuole = (The corrected fluorescence density of the target proteins localized at PM) / (The corrected fluorescence density). The ratios of GFP-Rsp5, Art5-GFP and Art1-mNG recruitment were measured from n=20 cells.

## Acknowledgements

We are grateful to Dr. Jason A. MacGurn, Dr. Matthew G. Baile, and Dr. Sho Suzuki for critical reading of the manuscript. We also thank other members of the Emr lab for helpful discussions. This work was supported by a Cornell University Research Grant (CU563704) to Scott D. Emr.

## Figure legends

**Figure S1.** Art1 and Art4 undergoes K63-linked di-ubiquitination. (A) Scheme of the Art1 domains. (B) Immunoblot analysis of Art1, *art1^K486R^*, *art1^ΔPY^*, *art1^ΔPY^*-1xUb, *art1^ΔPY^*-2xUb and *art1^ΔPY^*-3xUb in the wild-type strain. (C) Immunoblot analysis of Art5, *art5^K364R^*, *art5^ΔPY^* in both the *Ub-WT* and *Ub-K63R* mutant strains. (D) Art1 is di-ubiquitinated in a K63 linkage at the residue K486. (E) Art4 domain architecture. (F) Immunoblot analysis of Art4, *art4^K364R^*, *art4^ΔPY^*, as well as in *art4^ΔPY^*-1xUb, *art4^ΔPY^*-2xUb and *art4^ΔPY^*-3xUb in both the *Ub-WT* and *Ub-K63R* mutant strains. (G) Art4 is di-ubiquitinated in a K63 linkage. The whole cell lysate protein samples were resolved on 7% SDS-PAGE gels and the blot was probed with FLAG and GAPDH antibodies.

**Figure S2.** Ubiquitinated Art1 is required for efficient Mup1 ubiquitination. (A) Mup1 degradation in the yeast mutant *art1*Δ expressing empty vector, *tetO7*-Art1^WT^ or *tetO7*-Art1^K486R^. (B) Fluorescence microscopy of Mup1-GFP and Vph1-mCherry with or without methionine treatment. (C) Quantification of full length Mup1-GFP of the blots in (A). (D) IP of Mup1-GFP and Art1. (E) IP of Mup1-GFP and Art1^WT^ and *art1^Q49R^*. (F) IP of Mup1-GFP and Art1^WT^ and *art1^K486^.* (G) Western blot analysis of Art1^WT^ and *art1^K486^* in both WT and *npr1*Δ mutant. (H) Western blot analysis of Art1^WT^ and *art1^K486^* in WT cells with rapamycin (1µg/ml) or cycloheximide (50µg/ml) treatment for 1hour. (I) Cell growth assay of *art1*Δ mutant expressing *art1*^ΔPY^, *art1*^K486R^, Art1-1xUb, Art1-2xUb, Art1-3xUb, *art1*^K486R^-1xUb, *art1*^K486R^-2xUb or *art1*^K486R^-3xUb grown at 30°C for 3 days on synthetic media containing canavanine. (J) Cell growth assay of *art1*Δ mutant expressing *art1*^ΔPY^-1xUb, *art1*^ΔPY^-2xUb or *art1*^ΔPY^-3xUb grown in synthetic media with canavanine at 30°C for 3 days.

**Figure S3.** The Art1 di-ubiquitination facilitates Rsp5 PM recruitment upon methionine treatment. (A-C) Fluorescence microscopy of Art1-mNeonGreen (mNG) WT, K486R and PY motif mutants treated with methionine or shifted from minimal media to rich media for 1hr. (D) Quantification of Art1 recruited to PM (%) in the experiment of (A-C), (E) Localization of GFP-Rsp5 in the presence of Art1^WT^, *art1*^ΔPY^ or *art1*^K486R^. (F) Quantification of PM recruitment of Rsp5 in the experiment (E). (G) GFP-Rsp5 PM recruitment in the yeast cells expressing MUP1, *mup1*Δ or *mup1*-Q49R mutant. (H) Quantification of the Rsp5 PM recruitment in the experiment (G).

**Figure S4.** Deubiquitination of adaptor protein Art1 and Art5 by Ubp2. (A) Western blot analysis of Art1^WT^ and *art1^K486R^* mutant in the indicated yeast strains: RSP5(WT*), rsp5-*Y516A and *rsp5-* F618A. (B) Western blot analysis of Art1-HTF with overexpression of yeast DUBs proteins individually. (C) Western blot analysis of Art1-HTF in the *ubp2*Δ mutant bearing an empty vector, or with overexpression of UBP2 or *ubp2*(C745V) mutant. (D) Western blot analysis of Art1-HTF in yeast strains: RSP5(WT*), rsp5-*F618A, *ubp2*Δ and *rsp5-F618A*/*ubp2*Δ. (E) Western blot of Art1-HTF in *ubp2*Δ and *rsp5-F618A*/*ubp2*Δ yeast strains bearing an empty vector, UBP2 or *ubp2*(C745V) mutant. (F) Western blot analysis of tetO7-Art5-HTF in WT and *ubp2*Δ mutant with mock treatment or inositol treatment (1hr). After inositol treatment, cells were washed and grown in fresh media for 3 hours. (G) Western blot analysis of tetO7-Art1-HTF in WT and *ubp2*Δ mutant with or without methionine treatment (1hr). The methionine treated cells were then washed and grown in fresh media for 3hours. (H) Cartoon model depicting the Art1 is ubiquitinated by E3 ligase upon environmental cue then deubiquitinated by Ubp2. Non-ubiquitinated form of Art1 is ubiquitinated at K486 residue and engaged by Rsp5 for activation. This activated form of Art1 is then deubiquitinated by Ubp2 and Non-ubiquitinated form of Art1 is dis-engaged from Rsp5.

