## Supplementary figures and images for "Adaptor linked K63 di-Ubiquitin activates Nedd4/Rsp5 E3 ligase"

### Supplemental figures

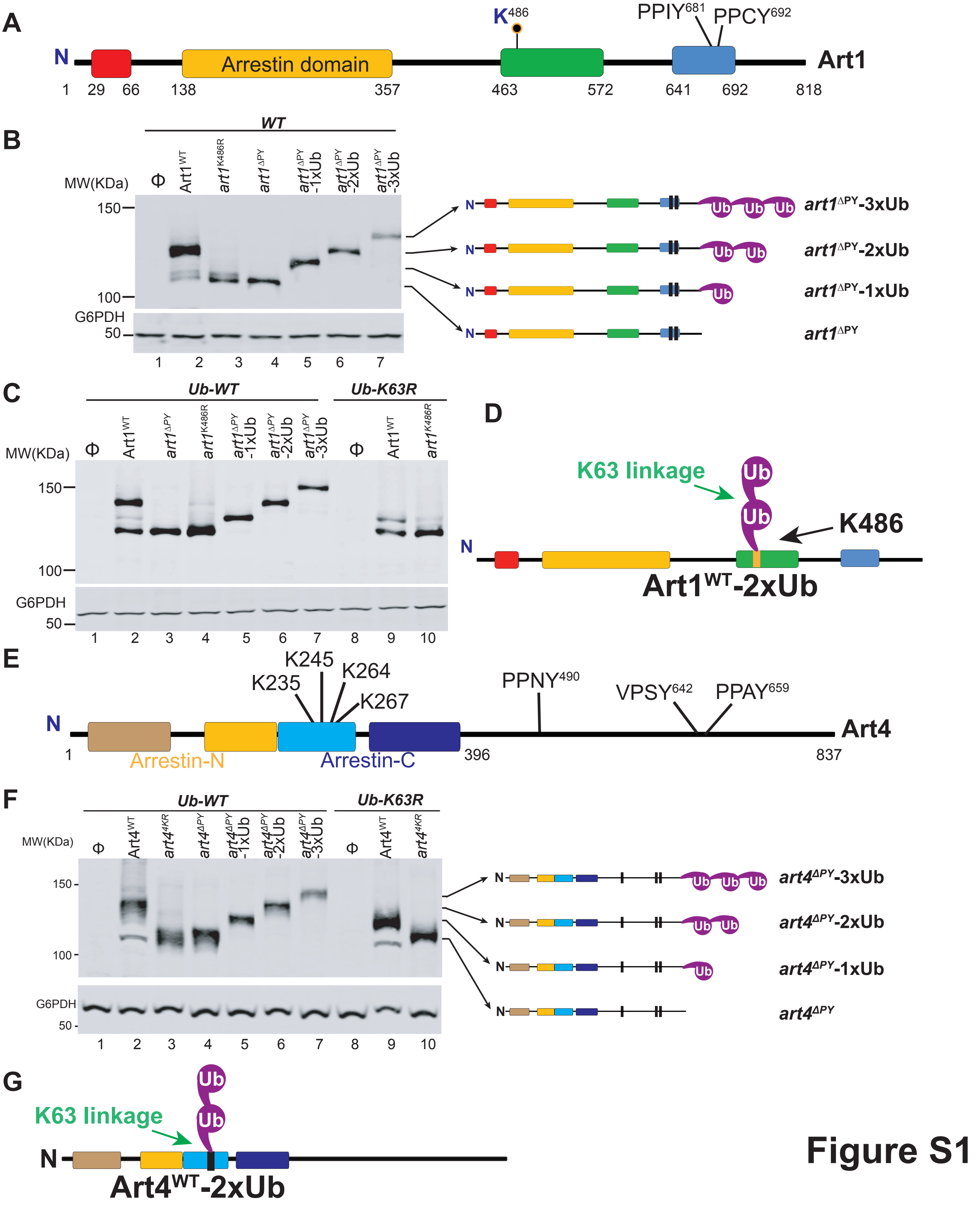

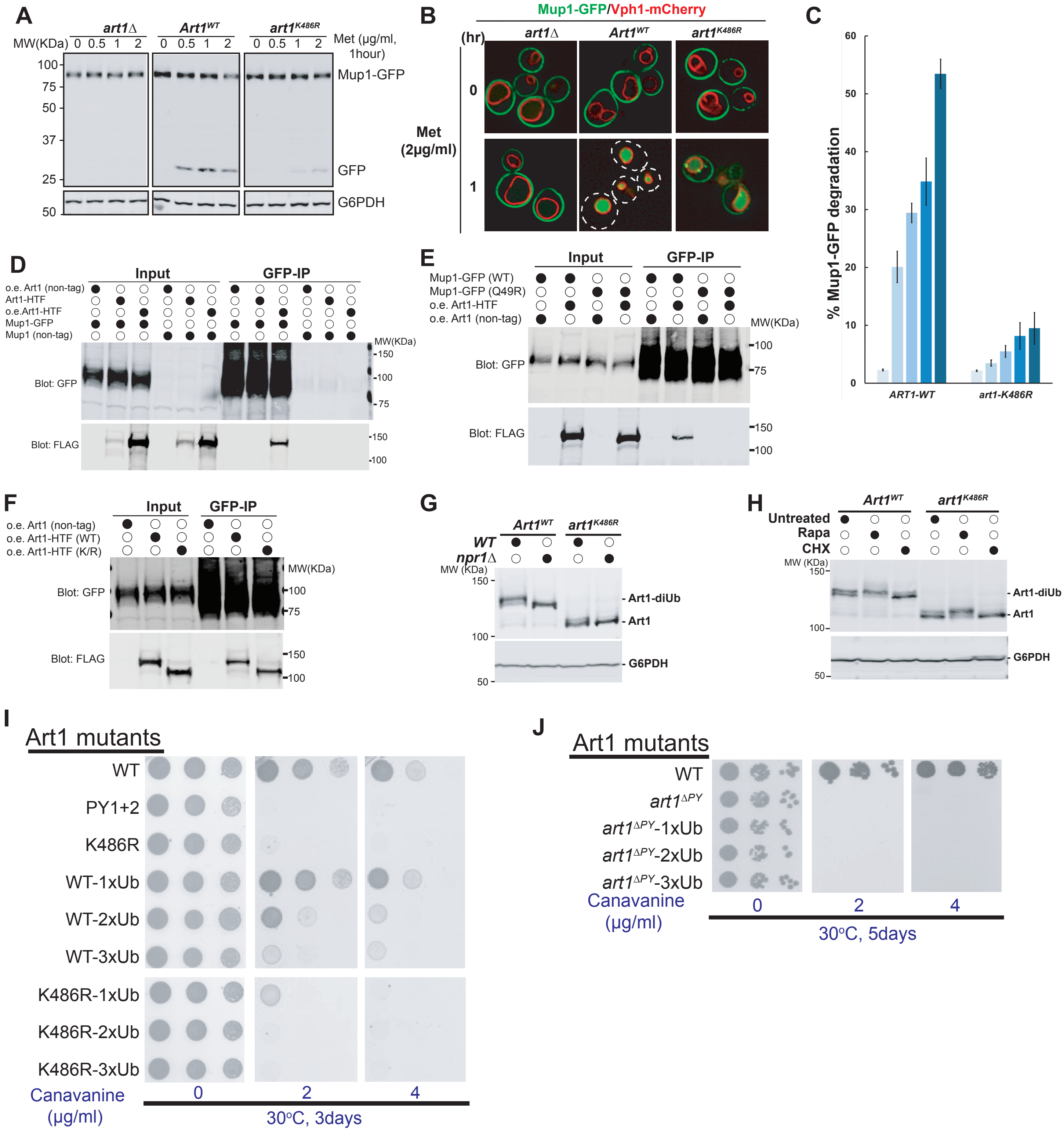

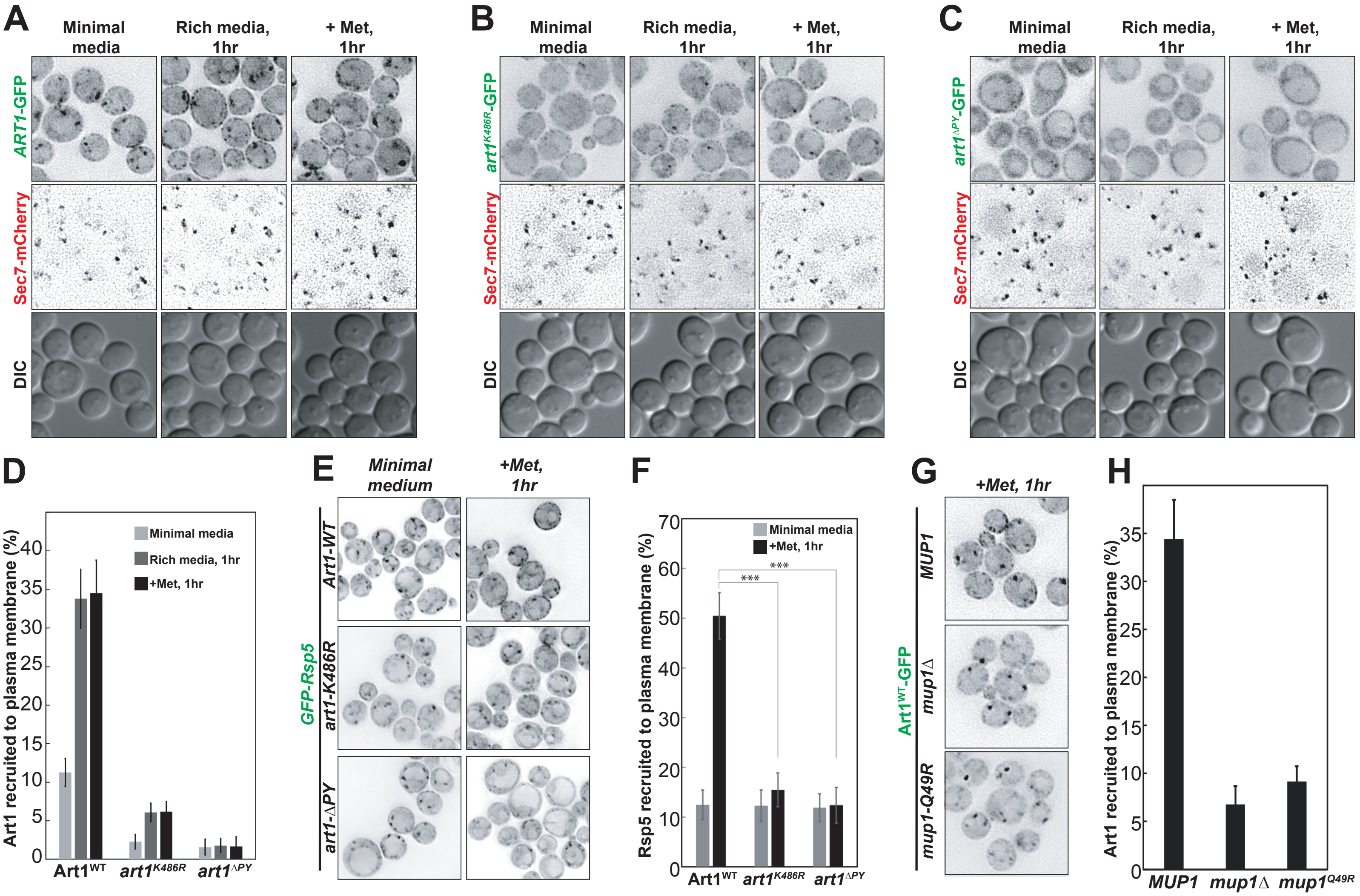

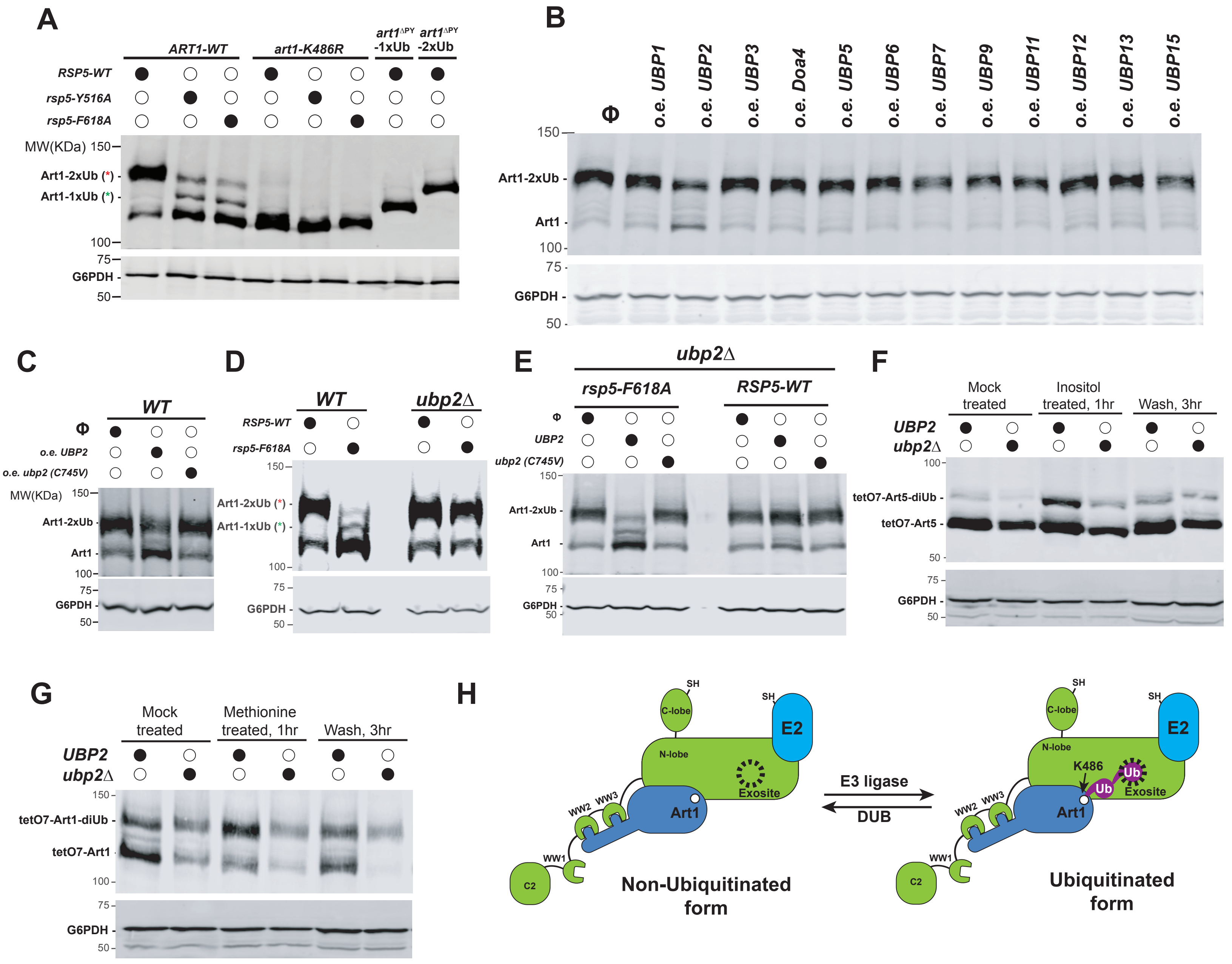
